# APOBEC-mediated DNA alterations: a possible new mechanism of carcinogenesis in EBV-positive gastric cancer

**DOI:** 10.1101/473884

**Authors:** Irina Bobrovnitchaia, Renan Valieris, Rodrigo Drummond, João Lima, Helano Freitas, Thais Bartelli, Maria Amorin, Diana Nunes, Emmanuel Dias-Neto, Israel Silva

**Affiliations:** A.C. Camargo Cancer Center

## Abstract

Mechanisms of viral oncogenesis are diverse and include the off-target activity of enzymes expressed by the infected cells, which evolved to target viral genomes for controlling their infection. Among these enzymes, the single-strand DNA editing capability of APOBECs represent a well-conserved viral infection response that can also cause untoward mutations in host DNA. Here we show, after evaluating somatic single-nucleotide variations and transcriptome data in 240 gastric cancer samples, a positive correlation between APOBEC3s mRNA-expression and the APOBEC-mutation signature, both increased in EBV+ tumors. The correlation was reinforced by the observation of APOBEC-mutations preferentially occuring in transcriptionally-active loci. The EBV-infection and APOBEC3 mutation-signature axis was confirmed in a validation cohort of 112 gastric cancer patients. Our findings suggest that APOBEC3 upregulation in EBV+ cancer may boost the mutation load, providing further clues to the mechanisms of EBV-induced gastric carcinogenesis. After further validation, this EBV-APOBEC axis may prove to be a secondary driving force in the mutational evolution of EBV+ gastric tumors, whose consequences in terms of prognosis and treatment implications should be vetted.

## 1. Introduction

Viruses have been implicated as etiologic agents in the development of human cancer, such as high-risk human papillomaviruses (HPVs), hepatitis B and C viruses (HBV, HCV), human T cell lymphotropic virus-1 (HTLV-1), the Kaposi’s sarcoma herpesvirus (KSHV), as well as the Epstein-Barr virus (EBV). Together these viruses are associated with 12-20% of human cancers worldwide [1,2] and further evidence suggest a possible link between viruses and other tumor types [3]. As described before, the molecular mechanisms of viral oncogenesis involve complex multistep processes causing the activation of cancer-causing pathways, which further on lead to the acquisition of cancer hallmarks [1,2]. These mechanisms include the expression of a number of viral oncoproteins that directly contribute to the neoplastic transformation and, more indirectly, through inflammatory responses triggered by chronic infections that may lead to lifelong persistent viral infection [1].

Nevertheless, despite the fast mutation and evolution rates of particular viruses, a mutual adaptation between the host and the infectious agents takes place, including the reduced replication and expression of viral proteins, the modulation of the host response by viral-miRNAs, as well as the emergence of innate and adaptive immunity [4, 5, 6, 7, 8]. In addition to these mechanisms, cytokines produced by the host as a response to viral infections induce the expression of AID/APOBEC (Activation-Induced cytidine Deaminase/Apolipoprotein B Editing Complex) family genes, which are important for antibody maturation and inhibition of viral infections through mutagenic and non-mutagenic mechanisms. The increased expression of some members of the APOBEC3 is one of the mechanisms used to control viral infections, as well as the retrotransposition of endogenous retroelements through the deamination of cytosines to uracil in single-stranded DNA (ssDNA) at replication forks, transcriptionally active genomic bubbles or RNA [9,10, 11].

These mechanisms are efficient in disrupting viral DNA replication and transcription, but an important side effect is the accumulation of off-target genomic mutations in the host DNA. The APOBEC-induced DNA editing does not occur randomly. Instead, it takes place in very specific DNA motifs, T<u>C</u>W→<u>T</u>W or T<u>C</u>W→T<u>G</u>W (mutated nucleotide underlined, W=A or T). These patterns are known as APOBEC mutation signatures [11].

In spite of it’s specificity, normal APOBEC activity has an intrinsic off-target deaminase activity over human cellular RNA and DNA [12]. This will lead to the accumulation of genomic mutations in the host DNA and also poses a threat to the integrity of the host genome and can itself become oncogenic. Therefore APOBEC may be especially active in cancers related to viral infection, being a second driving force towards to neoplastic transformation. Here we assessed the relationship of EBV infection, the expression of APOBEC3 genes and APOBEC mutation hallmarks in the gastric cancer (GC) TCGA cohort [13], followed by the validation of the EBV-APOBEC mutational signature in an independent cohort, in an attempt to investigate whether the presence of EBV-infection would trigger APOBEC-dependent DNA damage.

## 2. Results

### 2.1 The mRNA expression of APOBEC3s varies according to EBV status in GC samples

The expression levels of APOBEC3 genes were recovered from the total TCGA GC cohort, using the original pipelines of this project (see Methods section). We considered all cases for which EBV-status was informed and RNA-Seq and exome data were available [13], comprising a total of 240 subjects. We abode by the original TCGA four-tiered molecular classification: chromosomal instability (CIN, N: 122); microsatellite instability (MSI, N:47); genomically stable (GS, N:48) and positive for EBV infection (EBV, N:23).

The results observed are consistent with a physiological response to viral infection as 6 out of 7 APOBECs genes showed to be upregulated in the 23 EBV-positive samples, in stark contrast to the set of 217 EBV-negative samples (Figure 1). With the single exception of APOBEC3A, the APOBEC3 gene with the lowest expression levels for all sample groups, all other APOBEC3s showed to be upregulated in the EBV-positive group compared EBV-negative samples. APOBEC3C was the most abundantly expressed APOBEC3 transcript among all sample groups.

**Figure 1:**
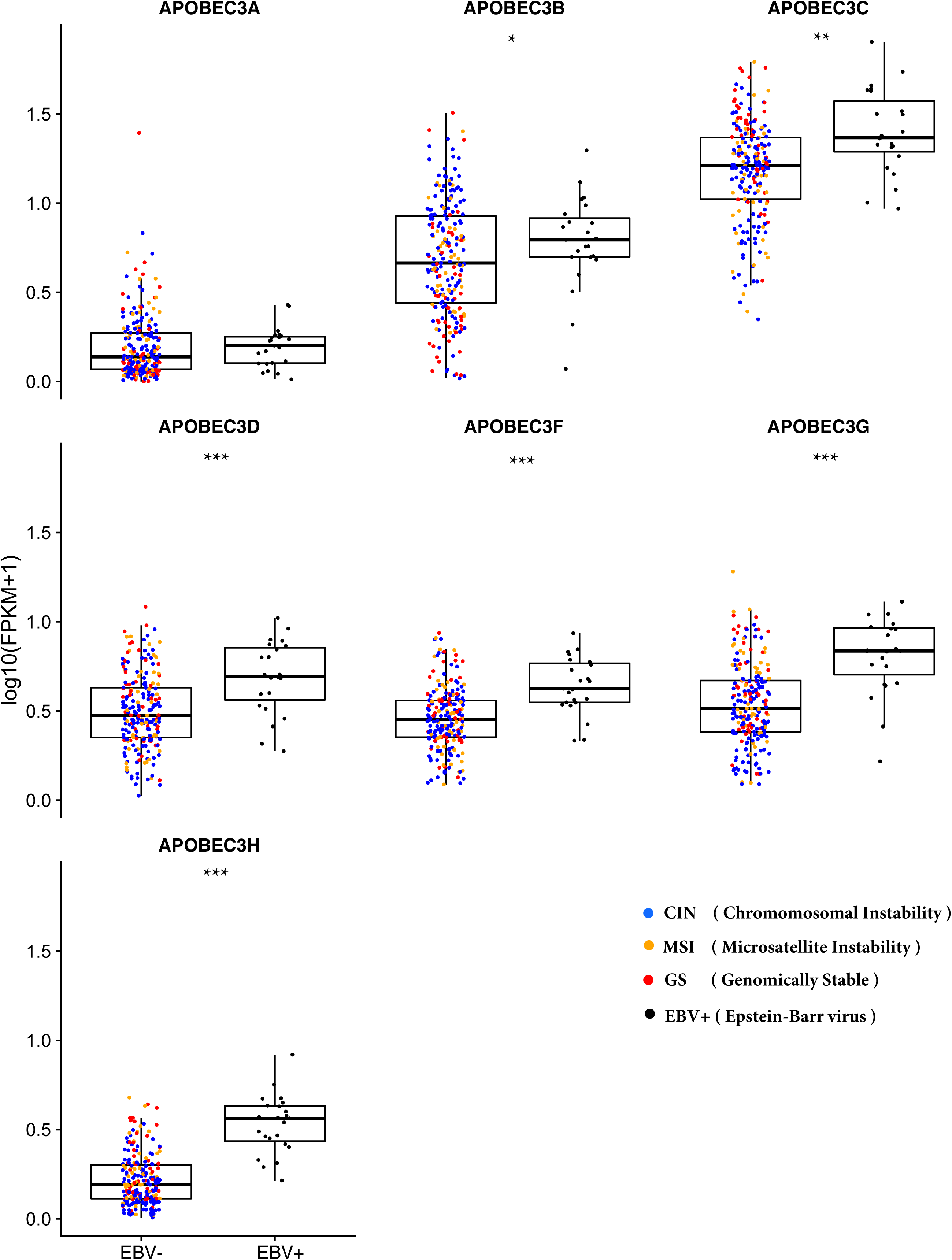
Comparison of expression levels of APOBEC3 members between EBV-negative (EBV-) and EBV-positive (EBV+) GC tumor types. GC-subtypes are indicated by colored dots as follows: MSI, Orange; GS, red; CIN, blue and EBV, black. Resulting p-values: *<=0.05; **<=0.0001 and ***<=0.00001 by Wilcoxon test.

### 2.2 EBV+ GC harbor a high load of APOBEC-mediated mutations

Next, for this same TCGA cohort, we evaluated the APOBEC mutation enrichment in EBV-negative and positive GC patients he TCW mutation enrichment [14] was used as a measure of the incidence of the APOBEC mutagenesis pattern. The TCW mutation pattern showed to be enriched in the EBV-positive as compared to EBV-negative samples (p = 0.01). This difference was validated an independent validation cohort of Brazilian GC patients (positive and negative for EBV) subjected to mutation analysis in a panel of 99 genes (p=0.042).

Having confirmed that TCW-mutation is enriched in EBV-positive as compared to EBV-negative tumors, we set out to investigate whether the increment in TCW-enrichment would correlate to an increment of APOBEC mRNA expression for the EBV-positive samples. For this, APOBEC3 expression was quantified by mapping RNA-Seq reads to each one of the APOBEC3 genes using the FPKM method (Section 2.1 and Methods). These expression levels were plotted against the enrichment for the TCW mutation context for each sample, as described [14].

Interestingly, for six out of the seven known human APOBEC3 genes (86%), we found a positive correlation between TCW-mutation enrichment and APOBEC3 expression (Figure 3, left panel). In contrast, by extending this analysis to EBV-negative tumors, we observed no correlations between APOBEC3s expression and the mutation load in the TCW context (Figure 3, right panel). To highlight the most correlated APOBECs genes throughout the tumor subtypes, we performed a clustering analysis that showed mutations in TCW context to positively correlate with APOBEC transcription levels in EBV-positive samples (Figure 4).

**Figure 2:**
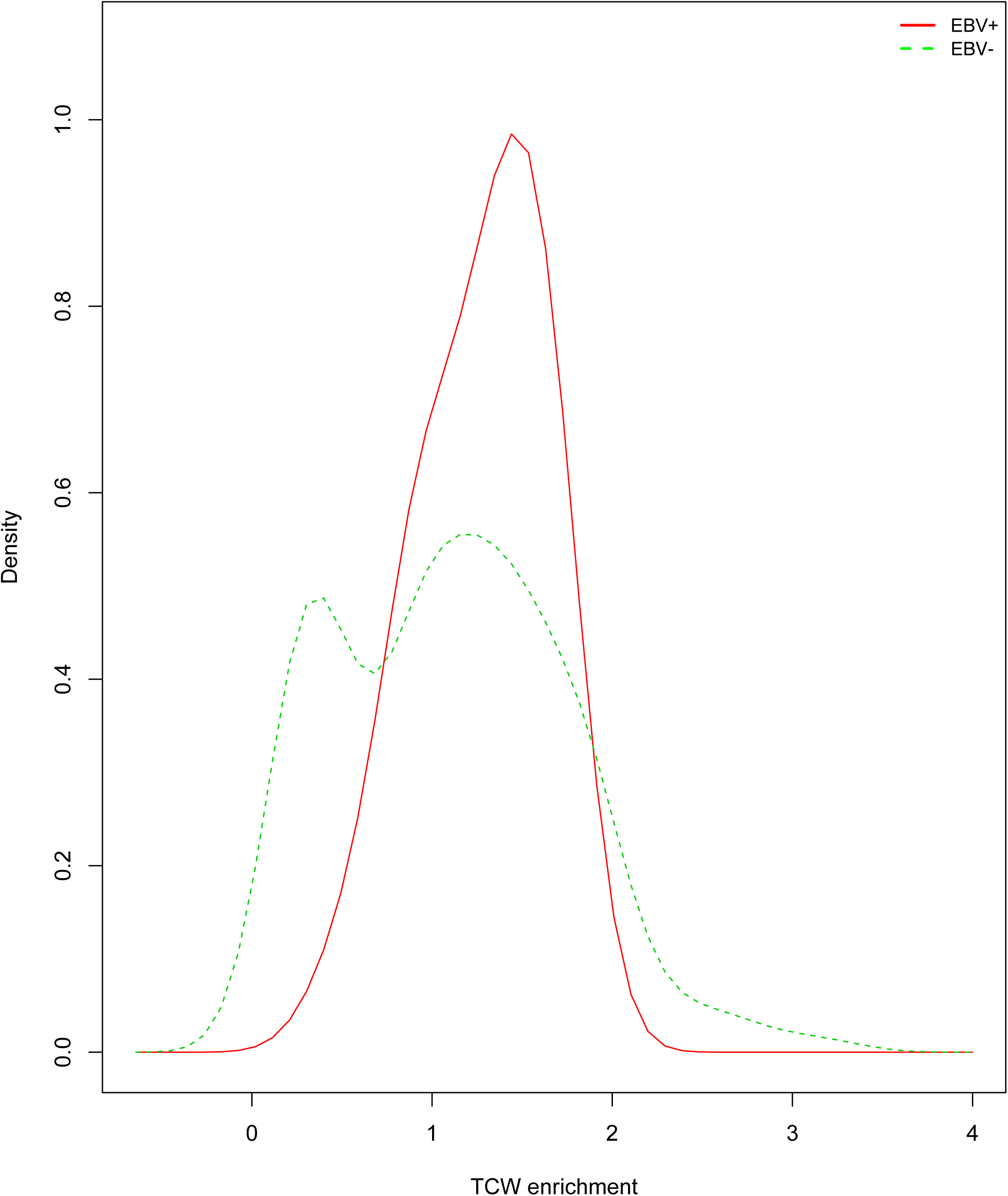
Analysis of TCW enrichment density between EBV-positive and EBV-negative samples. A permutation test of equality between the two densities indicates that the distributions are significantly different (p=0.01).

**Figure 3:**
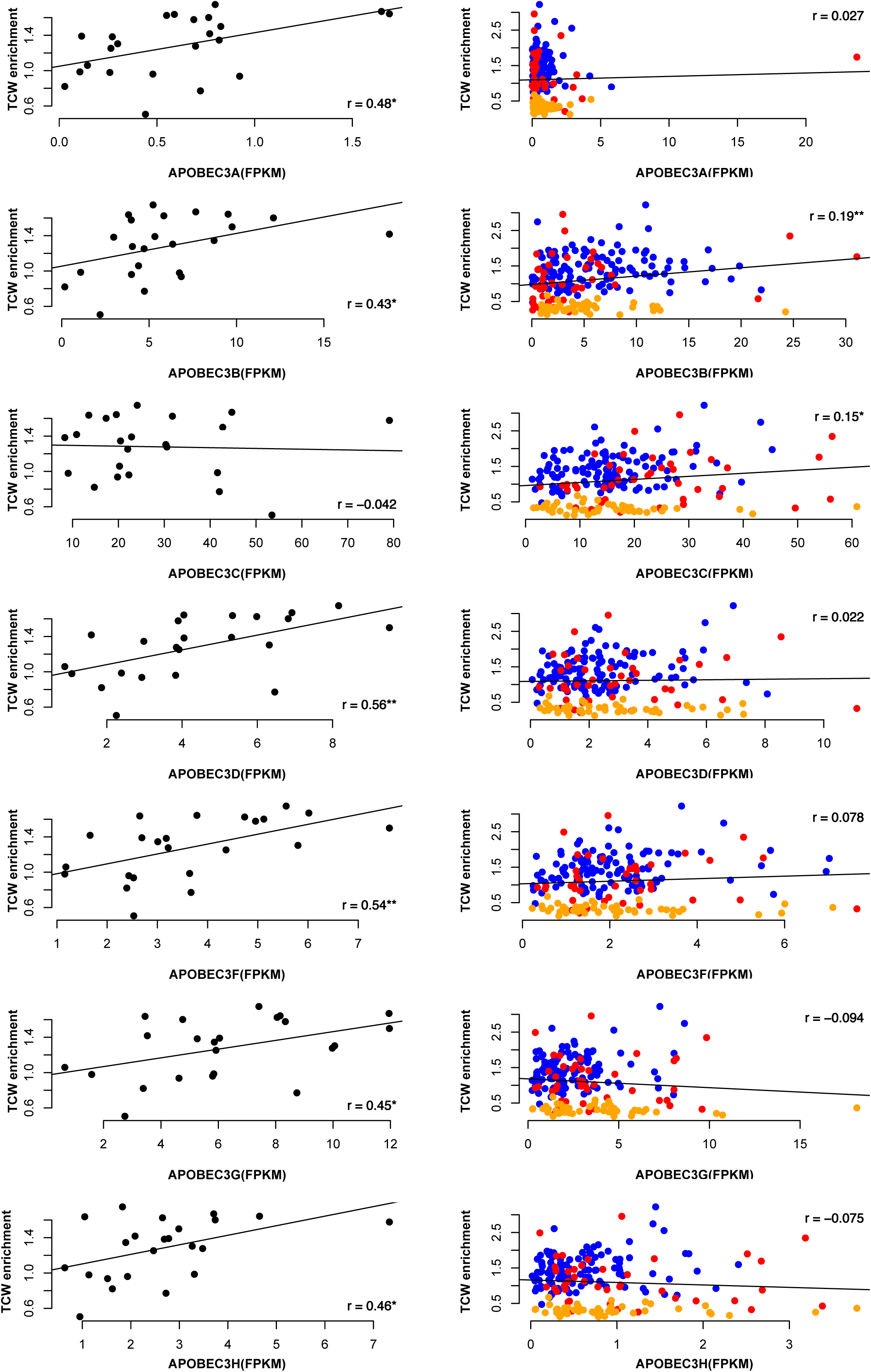
Comparison of expression levels of APOBEC3 genes and APOBEC3-related mutations (TCW enrichment). The scatter plot shows that elevated expression of most of the APOBEC3 genes is associated with higher APOBEC3-related mutations in EBV-positive samples. GC-subtypes are indicated by colored dots as follows: MSI, Orange; GS, red; CIN, blue and EBV, black. Resulting p-values: *<=0.05; **<=0.01 by test for association between paired samples.

**Figure 4:**
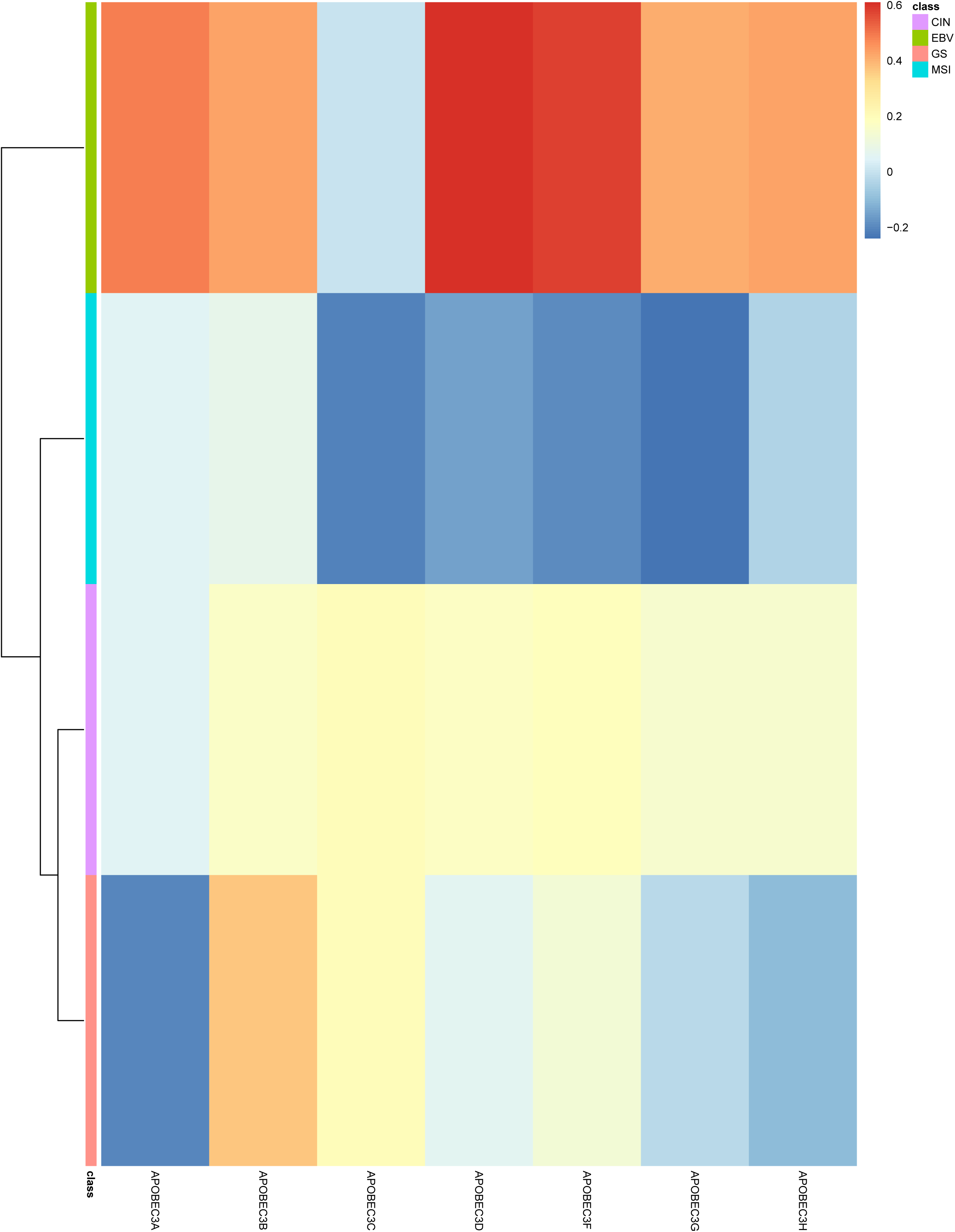
Heatmap for all correlations between gene expression levels of APOBEC3-family members and the TCW enrichment of APOBEC3-related mutations throughout the tumor subtypes. The gene expression values for the APOBEC3-family members are given as FPKM.

In order to determine if the APOBEC activity detected by the RNA-seq reads, would likely derive from the tumor cells or from other cells present in the tumor sample, we evaluated a possible correlations between APOBEC gene expression and tumor purity, using the tumor purity score given by TCGA. The results showed that APOBEC expression positively correlates with tumor purity, a finding that clearly suggests that APOBEC activity is indeed derived from the tumor cells (Figure 5).

**Figure 5:**
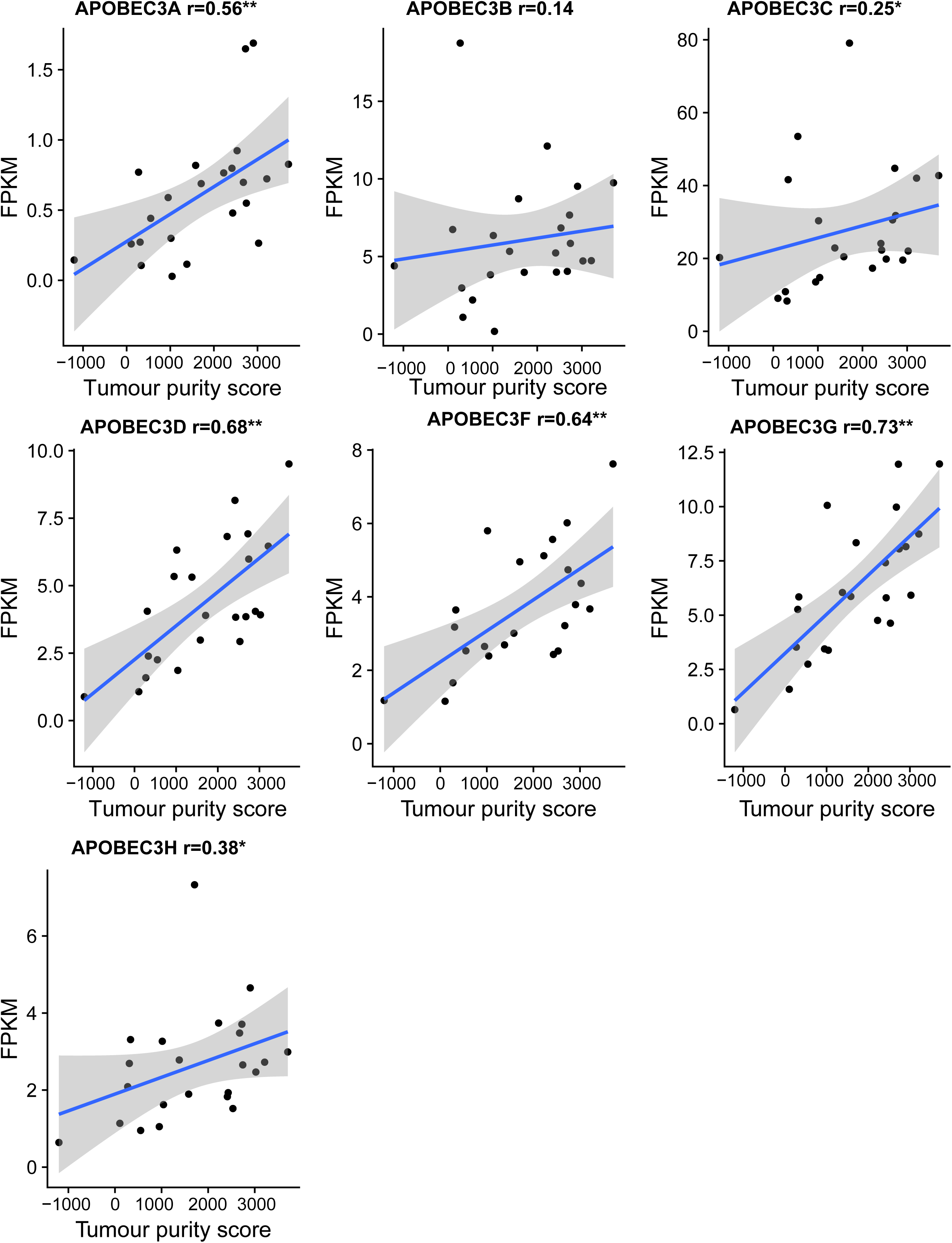
Correlation between tumor purity and APOBECs gene expression. Resulting p-values: *<=0.05; **<=0.0001 by test for association between paired samples.

### 2.3 APOBEC mutations in transcriptionally active genes

Since the increased TCW motif mutation load seen in EBV-positive samples is indeed correlated with the higher mRNA expression of APOBEC3 genes, we conceived that the APOBEC3 mutational load would be correlated to the expression of the mutated genes; i.e. highly expressed genes should display more mutations than genes with lower expression, due to greater availability of transcriptionally active ssDNA bubbles, the preferential editing sites of APOBEC3s. In order to estimate this, we divided all transcripts detected in the TCGA gastric cancer samples into 5 categories of transcriptional activity quantiles (see Methods) and evaluated the frequency of TCW-mutations in each one of these categories. In agreement with previous descriptions of an enhanced mutational load in ssDNA strands exposed during transcription, we found a clear trend for an increased load of TCW-motif alterations that parallels the expression levels of the transcripts in the different abundance categories. Indeed, a parallel of expression-mutation is clearly observed. In contrast, no trends were observed for the EBV-negative samples (Figure 6). Altogether, our findings indicate that whereas APOBEC3-mediated mutations damage a series of distinct transcripts, there is an enrichment on the mutational load in genomic regions associated with elevated transcriptional activity.

**Figure 6:**
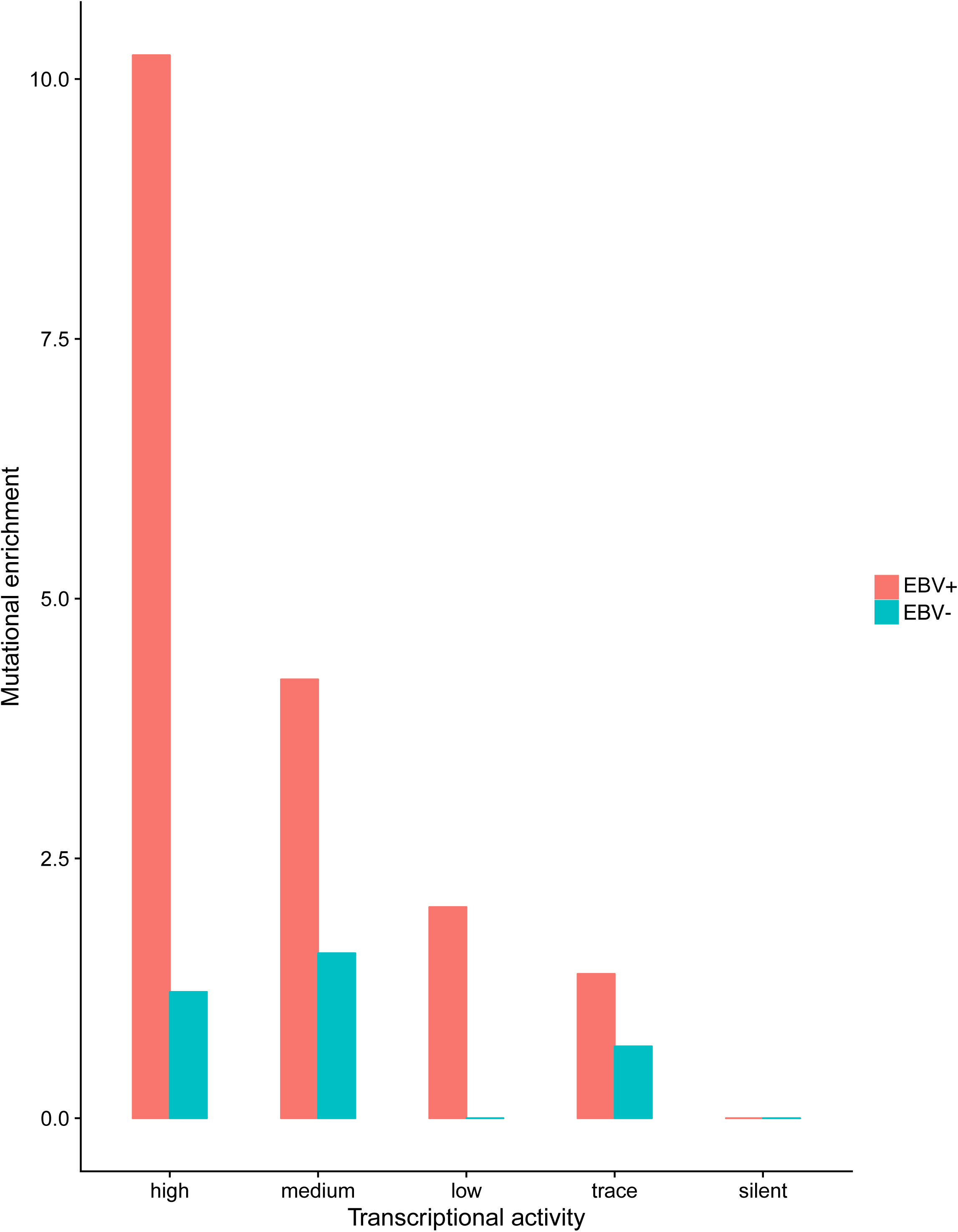
The mutation load in TCW context was found to be proportional to the transcriptional levels of the distinct genes in EBV-positive samples.

## 3. Discussion

The mechanisms of tumorigenesis so far described for EBV-induced tumors include the interference of EBV with cell-cycle checkpoints (G1 as well as G2/M transitions and the assembly of the mitotic spindle), the inactivation of cell death pathways, when immortalized EBV-infected B-cells proliferate in a central neoplastic mechanism of B-cell lymphomas [15] as well as the genome-wide reactivation of enhancers and promoters located near cryptic and otherwise repressed long terminal repeats of inactive human endogenous retroviruses [16]. In these cells, EBV-infection acts as a somatic mutator, leading to the activation of AID, an enzyme that mediates class switch recombination of immunoglobulin genes and induces cytosine deamination and somatic hypermutation [17]. These findings indicate that AID has a role in EBV-induced lymphomagenesis in B cells.

Here we investigated whether EBV-infection would trigger collateral APOBEC-mediated damage of the host genome in EBV-positive gastric cancer. We performed a comprehensive analysis in a large number of paired exome and expression profiles of GC samples derived from The Cancer Genome Atlas (TCGA) database. We found the expression of APOBEC3-family genes to be significantly increased in EBV-positive compared to EBV-negative gastric cancer. An intriguing study also found the presence of EBV DNA in breast cancer samples and was able to correlate an EBV-related transcriptional signature to the APOBEC mutation signature [18]. Furthermore, we showed a consistent correlation between the expression of six out of the seven APOBEC3 genes and mutations driven by APOBEC3 in EBV-infected gastric cancer. In contrast, EBV-negative tumors had no APOBEC3 overexpression nor relevant APOBEC-mediated mutations.

Whereas it has been described that EBV infection can induce the infiltration of immune cells [19] and we can not rule out the importance of APOBEC expression in immune infiltrating cells, we should mention that according to TCGA data, the percentage of tumor cells in the samples used here did not show significant differences among GC molecular subtypes - i.e. EBV+ and others - making less likely that the evaluated samples have significant discrepancies in terms of infiltrating immune cells [13]. Moreover, our correlation analysis between tumour purity and APOBEC gene expression showed that APOBEC activity is likely to be indeed derived from the tumor cells.

As we observed increased rates of TCW-context mutations at highly active transcriptional genomic loci, the presence of mutations apparently associated with exposed ssDNA may serve as a possible source of genomic instability in EBV-positive cancers. Taken together, our findings suggest that the observed antiviral APOBEC3 upregulation response might boost the mutation rates during viral carcinogenesis in EBV-positive gastric cancer. In this context, APOBEC activation may be a secondary mutation mechanism in EBV-related gastric cancers. These results gained support from our validation cohort that corroborated the correlation between EBV-status and TCW-mutation patterns. However, as the frequency of EBV-positive GC cases is low, larger cohorts and experiments with animal models are needed to further validate our findings. Nevertheless, APOBEC3s may emerge as a secondary driving force in the mutational evolution of EBV-positive gastric cancer and should have its consequences assessed in terms of prognosis, generation of neoantigens, immunotherapy and other therapeutic implications.

## 4. Materials and Methods

### 4.1 Genomic and clinical data in the Discovery Cohort

Somatic Single Nucleotide Variations (SNVs), clinical data as well as the normalized gene expression levels (FPKM, Fragments Per Kilobase of exon per Million fragments mapped) of 240 gastric cancer cases were downloaded from The Cancer Genome Atlas (TCGA) [13], and used as the discovery cohort. SNVs were processed through the internal pipeline at Genomic Data Commons (GDC) as tumor-normal pairs, which applies three separate somatic variant calling algorithms. We used a variant set detected by Mutect2 which employs a "Panel of Normals" to filter out additional germline mutations. These datasets derive from four different tumor subtypes, as follows: Epstein–Barr virus (EBV+, N=23), microsatellite instability (MSI, N=47), genomically stable (GS, N=48) and chromosomal instability (CIN, N=122). Characteristics of participants in the discovery cohort are available to all interested upon request.

### 4.2 Validation Cohort: Library preparation, NGS and EBV analysis

The study was approved by the Institutional Review Board of the A.C.Camargo Cancer Center (protocol 2134/15), São Paulo, Brazil. All 112 GC patients of the validation cohort were prospectively enrolled in an institutional study to unveil the epidemiology and genomics of gastric adenocarcinomas in Brazil [20]. All participants provided written informed consent either for this study or for the institutional tumor biobank.

With the exception of one case (1/112), all other surgical specimens were collected before any interventions and were provided by the institutional biobank. Genomic DNA (100ng) from gastric adenocarcinoma samples was used for capture-based enrichment of 781.132 bp of a customized gene panel including 99 of the most frequently mutated genes in gastric cancer, according to TCGA and to the literature. Libraries were prepared following the manufacturer’s instructions (SeqCap EZ Library SR User’s Guide Version 5.1 - NimbleGen, Roche), and were sequenced in the NextSeq 500 platform (Illumina), using paired-end reads (2×75bp).

The EBV genotyping was performed for all samples, as follows. Samples were analyzed by Real Time PCR with primers directed a 84-bp fragment of the BamH1W region of the EBV genome (F: 5'-GCAGCCGCCCAGTCTCT-3'; R: 5'-ACAGACAGTGCACAGGAGCCT-3') together with a labelled internal probe (FAM-AAAAGCTGGCGCCCTTGCCTG-TAMRA) [21,22] EBV detection was done together with an internal control directed to the amplification of a fragment of the human β-actin gene (F: 5′- CCATCTACGAGGGGTATGC-3′; R: 5′-GGTGAGGATCTTCATGAGGTA-3′), together with an internal probe (VIC-CCTGCGTCTGGACCTGGCTG-NFQ), in a multiplex reaction. Amplicons were detected in real time in an ABI 7500 Fast Real Time instrument. Thermocycling conditions were: 95°C for 20 seconds, 95°C for 3 seconds and 60°C for 30 seconds for 40 cycles. Each 20ul reaction contained: TaqMan^®^ Fast Universal Master Mix (2x) (Applied Biosystems^®^), as well as primers and probes (primers: 18µM; probes: 5µM) for each target.

To check for amplicon contamination, every run contained at least two “no template” controls (water). Finally, the determination of EBV infection status was performed by the ΔCt method, where EBV Ct was subtracted from β-actin Ct, and the result calculated by 2^(-ΔCt). the samples were considered positive for EBV infection when the ratio EBV/ACTB was of at least 1.

### 4.3 SNV calling in the Validation Cohort

For the validation cohort, somatic SNVs were called through an in-house pipeline, following the Broad Institutes GATK best practices [23]. Briefly, raw reads were aligned using BWA-mem (Burrows Wheeler Aligner) [24] to assembly hg19/GRCh37. Alignment files in SAM format were converted to BAM files, sorted and then filtered to exclude reads with mapq score < 15. Retained reads were processed using SAMtools [25] and Picard (https://broadinstitute.github.io/picard/) respectivaly, which excludes low-quality reads and PCR duplicates. Finally, somatic SNVs call was performed for the panel data from analysis-ready BAM files using GATK-UnifiedGenotyper (v3.8) for tumor samples and filtered with a panel of 16 unmatched blood samples. Extensive filtering was applied to remove low mapping quality, as well as strand and position bias. Further residual germline variants were filtered out using the the database of germline mutations of the Exome Aggregation Consortium (ExAC) [26] and the Online Archive of Brazilian Mutations (ABraOM) [27].

### 4.4 Detecting an APOBEC3 mutation enrichment

We elected the stringent TCW motif (where W = A or T) as the preferred sequence context for APOBEC3-mediated cytosine deamination [11]. The enrichment score in the TCW-context was considered as a measurement of prevalence of APOBEC mutation (as described by [14]) and calculated as

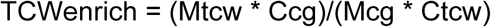

where Mtcw is the number of mutated cytosines/guanines located in a TCW motif, Mcg is the total number of mutated cytosines/guanines, Ctcw is the total number of TCW within the +/-20 nucleotides window around mutated cytosines/guanines and Ccg is the total number of cytosines/guanines within the +/-20 nucleotides window around mutated cytosines/guanines.

### 4.3 Mutation enrichment in expression genes levels

We evaluated the mRNA expression levels for all seven APOBEC3-family genes: *APOBEC3A, APOBEC3B, APOBEC3C, APOBEC3D, APOBEC3F, APOBEC3G,* and *APOBEC3H* in all gastric cancer samples. For this we considered the gene expression levels processed and provided by the TCGA pipelines, from their public database (https://portal.gdc.cancer.gov/).

The quantitative expression levels were obtained by counting the mapped sequence reads, normalized through the FPKM method (Fragments Per Kilobase of exon per Million fragments mapped) [28]. This approach considers the uniquely mapped reads of the RNA-Seq experiment (depth of sequencing), length of the transcripts, as well as the number of counts for all exons of each transcript. Next, after separating the transcripts with no expression (silent) we stratified the active transcripts in four levels (high, the top 25%; medium, 75 to 50%; low, 50 to 25% and trace, those below 25%) based on FPKM values using a normal mixture model implemented in the Mclust and quantile methods employed in R language as previously described [29]. Then we calculated a numeric value to evaluate the APOBEC-induced mutations in those classes for each gastric cancer sample as follows:

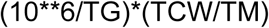

where TG is the total of genes with any mutation, TCW is the total number of mutations in APOBEC-context for each class and, finally, TM is the total of mutations for each class.

### 4.4 Association with tumor purity

In addition to the tumor cells, tumor microenvironment typically comprises a mixed population of cells, including stromal and infiltrating immune cells. To evaluate whether tumor purity (percent tumor cells) correlates with APOBEC expression, we took advantage of the tumor purity inferred from ESTIMATE (Estimation of STromal and Immune cells in MAlignant Tumor tissues using Expression data) [30] available at MD Anderson Cancer Center (https://bioinformatics.mdanderson.org/estimate/). We used Pearson’s correlation coefficient as a test for association between tumor purity and gene expression.

### 4.5 Statistical Analysis

The comparison of gene expression and TCW enrichment between the EBV-positive and EBV-negative samples were performed using the nonparametric Wilcoxon's signed rank test (one-tailed unpaired test) or a permutation test of equality as appropriate.

## References

1. White MK, Pagano JS, Khalili K. Viruses and human cancers: a long road of discovery of molecular paradigms. Clin Microbiol Rev. 2014 Jul;27(3):463-81.

2. Luo GG, Ou JH. Oncogenic viruses and cancer. Virol Sin. 2015. Apr;30(2):83-4.

3. Nüssing S, Sant S, Koutsakos M, Subbarao K, Nguyen THO, Kedzierska K. Innate and adaptive T cells in influenza disease. Front Med. 2018 Feb;12(1):34-47.

4. Teixeira AA, Marchiò S, Dias-Neto E, Nunes DN, da Silva IT, Chackerian B, Barry M, Lauer RC, Giordano RJ, Sidman RL, Wheeler CM, Cavenee WK, Pasqualini R, Arap W. Going viral? Linking the etiology of human prostate cancer to the PCA3 long noncoding RNA and oncogenic viruses. EMBO Mol Med. 2017 Oct;9(10):1327-1330.

5. Saez-Cirion A, Manel N. Immune Responses to Retroviruses. Annu Rev Immunol. 2018 Jan 12.

6. Münz C. Epstein-Barr Virus-Specific Immune Control by Innate Lymphocytes. Front Immunol. 218 2017 Nov 24;8:1658.

7. Albanese M, Tagawa T, Buschle A, Hammerschmidt W. MicroRNAs of Epstein-Barr Virus Control Innate and Adaptive Antiviral Immunity. J Virol. 2017 Jul 27;91(16).

8. Refsland EW, Harris RS. The APOBEC3 family of retroelement restriction factors. Curr Top Microbiol Immunol. 2013;371:1-27.

9. Siriwardena SU, Chen K, Bhagwat AS. Functions and Malfunctions of Mammalian DNA-Cytosine 225 Deaminases. Chem Rev. 2016 Oct 26;116(20):12688-12710.

10. Seplyarskiy VB, Soldatov RA, Popadin KY, Antonarakis SE, Bazykin GA, Nikolaev SI. APOBEC-induced mutations in human cancers are strongly enriched on the lagging DNA strand during replication. Genome Res. 2016 Feb;26(2):174-82.

11. Roberts SA, Sterling J, Thompson C, Harris S, Mav D, Shah R, Klimczak LJ, Kryukov GV, Malc E, 230 Mieczkowski PA, Resnick MA, Gordenin DA. Clustered mutations in yeast and in human cancers can arise from damaged long single-strand DNA regions. Mol Cell. 2012 May 25;46(4):424-35.

12. Conticello SG. Creative deaminases, self-inflicted damage, and genome evolution. Ann N Y Acad Sci. 2012 Sep;1267:79-85.

13. Bass AJ, Thorsson V, Shmulevich I, Reynolds SM, Miller M, Cancer Genome Atlas Research Network. Comprehensive molecular characterization of gastric adenocarcinoma. Nature. 2014 Sep 11;513(7517):202-9.

14. Roberts SA, Lawrence MS, Klimczak LJ, Grimm SA, Fargo D, Stojanov P, Kiezun A, Kryukov GV, Carter SL, Saksena G, Harris S, Shah RR, Resnick MA, Getz G, Gordenin DA. An APOBEC cytidine deaminase mutagenesis pattern is widespread in human cancers. Nat Genet. 2013 Sep; 45(9):97.

15. O’Nions J, Allday MJ. Deregulation of the cell cycle by the Epstein-Barr virus. Adv Cancer Res. 241 2004;92:119-86.

16. Leung A, Trac C, Kato H, Costello KR, Chen Z, Natarajan R, Schones DE. LTRs activated by Epstein-Barr virus-induced transformation of B cells alter the transcriptome. Genome Res. 2018 Oct 31. pii: gr.233585.117.

17. Ayako Arai, Masato Horino, Ken-Ichi Imadome, Kohsuke Imai, Honami Komatsu, Ludan Wang, Go Matsuda, Kasumi Hamazaki, Morito Kurata, Tetsuya Kurosu, Shigeyoshi Fujiwara and Osamu Miura. Epstein–Barr Virus Induces Activation-Induced Cytidine Deaminase Expression in T or NK Cells Leading to Mutagenesis and Development of Lymphoma. Blood 2013 122:1765.

18. Hu H, Luo ML, Desmedt C, Nabavi S, Yadegarynia S, Hong A, Konstantinopoulos PA, Gabrielson E, Hines-Boykin R, Pihan G, Yuan X, Sotiriou C, Dittmer DP, Fingeroth JD, Wulf GM. Epstein-Barr Virus Infection of Mammary Epithelial Cells Promotes Malignant Transformation. EBioMedicine. 2016 Jul;9:148-160.

19. Ma C, Patel K, Singhi AD, Ren B, Zhu B, Shaikh F, Sun W. Programmed Death-Ligand Expression Is Common in Gastric Cancer Associated With Epstein-Barr Virus or Microsatellite Instability. Am J Surg Pathol. 2016 Nov;40(11):1496-1506.

20. Soares FA, Coimbra FJF, Pelosof AG, GE4GAC group, Dias-Neto E. Genomics and epidemiology for gastric adenocarcinomas. Applied Cancer Research. 2017, 37:7.

21. Khoddami M, Nadji SA, Dehghanian P, Vahdatinia M, Shamshiri AR.. Detection of Epstein-Barr Virus DNA in Langerhans Cell Histiocytosis. Jundishapur J Microbiol. 2015 Dec 26;8(12):e27219.

22. Fan H, Nicholls J, Chua D, Chan KH, Sham J, Lee S, Gulley ML. Laboratory markers of tumor burden in nasopharyngeal carcinoma: a comparison of viral load and serologic tests for Epstein-Barr virus. Int J Cancer. 2004 Dec 20;112(6):1036-41.

23. Van der Auwera GA, Carneiro MO, Hartl C, Poplin R, Del Angel G, Levy-Moonshine A, Jordan T, Shakir K, Roazen D, Thibault J, Banks E, Garimella KV, Altshuler D, Gabriel S, DePristo MA. From FastQ data to high confidence variant calls: the Genome Analysis Toolkit best practices pipeline. Curr Protoc Bioinformatics. 2013;43:11.10.1-33.

24. Li H, Durbin R. Fast and accurate long-read alignment with Burrows-Wheeler transform. Bioinformatics. 2010 Mar 1;26(5):589-95.

25. Li H, Handsaker B, Wysoker A, Fennell T, Ruan J, Homer N, Marth G, Abecasis G, Durbin R; 1000 Genome Project Data Processing Subgroup. The Sequence Alignment/Map format and SAMtools. Bioinformatics. 2009 Aug 15;25(16):2078-9

26. Lek M, Karczewski KJ, Minikel EV, Samocha KE, Banks E, Fennell T, Exome Aggregation Consortium. Analysis of protein-coding genetic variation in 60,706 humans. Nature. 2016 Aug 18;536 (7616):285-91.

27. Naslavsky MS, Yamamoto G, De Almeida TF, Ezquina SAM, Sunaga DY, Pho N, Bozoklian D, Sandberg TOM, Brito LA, Lazar M, Bernardo DV, Amaro E Jr, Duarte YAO, Lebrão ML, Passos-Bueno MR, Zatz M. Exomic variants of an elderly cohort of Brazilians in the ABraOM database. Hum Mutat. 2017 Jul;38(7):751-763.

28. Mortazavi A, Williams BA, McCue K, Schaeffer L, Wold B. “Mapping and quantifying mammalian transcriptomes by RNA-Seq“. Nature Methods. 5: 621-8.

29. Klein IA, Resch W, Jankovic M, Oliveira T, Yamane A, Nakahashi H, Di Virgilio M, Bothmer A, Nussenzweig A, Robbiani DF, Casellas R, and Nussenzweig MC. Translocation-Capture Sequencing Reveals the Extent and Nature of Chromosomal Rearrangements in B Lymphocytes. Cell. 2011 Sep 30; 147(1): 95– 106.

30. Yoshihara K, Shahmoradgoli M, Martínez E, Vegesna R, Kim H, Torres-Garcia W, Treviño V, Shen H, Laird PW, Levine DA, Carter SL, Getz G, Stemke-Hale K, Mills GB, Verhaak RG. Inferring tumour purity and stromal and immune cell admixture from expression data. Nat Commun. 2013; 4():2612.

